# MXRA8 promotes adipose tissue whitening to drive obesity

**DOI:** 10.1101/2024.01.31.578211

**Authors:** Wentong Jia, Rocky Giwa, John R. Moley, Gordon I. Smith, Max C. Petersen, Rachael L Field, Omar Abousaway, Arthur S. Kim, Sarah R. Coffey, Stella Varnum, Jasmine M. Wright, Xinya Zhang, Samantha Krysa, Irfan J. Lodhi, Nada A. Abumrad, Samuel Klein, Michael S. Diamond, Jonathan R. Brestoff

**Affiliations:** Department of Pathology and Immunology, Washington University School of Medicine, St Louis, MO 63110, USA; Center for Human Nutrition, Washington University School of Medicine, St Louis, MO, 63110, USA; Department of Medicine, Washington University School of Medicine, St Louis, MO, 63110, USA; Department of Cell Biology and Physiology, Washington University School of Medicine, St. Louis, MO, 63110, USA; Department of Molecular Microbiology, Washington University School of Medicine, St Louis, MO, 63110, USA

**Keywords:** MXRA8, obesity, beige adipocytes, brown adipocytes, adipocyte progenitor cells, adipocyte, thermogenesis, UCP1

## Abstract

Matrix-remodeling associated 8 (MXRA8), also known as Dual immunoglobulin domain cell adhesion molecule (DICAM), is a type 1 transmembrane protein that reportedly binds the α_V_β_3_ integrin^1^ and regulates the differentiation of osteoclasts^2^ and chondrocytes^3^, tumor growth^4^, T cell trafficking^5^, and angiogenesis^6^. MXRA8 is also an essential entry receptor for chikungunya virus and other related arthritogenic alphaviruses.^7-9^ We compared MXRA8 expression in 51 tissues in the Human Protein Atlas and found it is most highly expressed in white adipose tissue (WAT), however the function of MXRA8 in WAT is unknown. Here, we found that MXRA8 expression in WAT is increased in people with obesity and that this response is also observed in a mouse model of high fat-diet (HFD)-induced obesity. Single-nucleus RNA sequencing and high-dimensional spectral flow cytometry analyses revealed that MXRA8 is expressed predominantly by adipocyte progenitor (AP) cells and mature adipocytes. MXRA8 mutant primary adipocytes from inguinal (i)WAT exhibited increased expression of Uncoupling protein 1 (UCP1), a thermogenic protein expressed by beige and brown adipocytes that limits obesity pathogenesis.^10-12^ Indeed, MXRA8 mutant mice fed a HFD had preserved UCP1^+^ beige and brown adipocytes and were protected from HFD-induced obesity in a UCP1-dependent manner. Collectively, these findings indicate that MXRA8 promotes whitening of beige and brown adipose tissues to drive obesity pathogenesis and identify MXRA8 as a possible therapeutic target to treat obesity and associated metabolic diseases.

## MAIN TEXT

To identify organ systems where MXRA8 is most highly expressed, we first examined the Human Protein Atlas (HPA, **Extended Data Fig 1a**) and Genotype-Tissue Expression (GTEx, **Extended Data Fig 1b**) datasets and found that MXRA8 expression in WAT was ranked 1^st^ of 51 tissues and 2^nd^ of 27 tissues, respectively. Based on this finding, we compared *MXRA8* transcript levels in subcutaneous abdominal white adipose tissue (WAT) from people who were metabolically healthy lean (MHL, n=15), metabolically healthy obese (MHO, n=18), or metabolically unhealthy obese (MUO, n=19), as defined previously.^13^ *MXRA8* expression was significantly increased in both obese groups compared with the metabolically healthy lean group and was highest in the metabolically unhealthy obese group (**Fig 1a**). The increase in WAT *MXRA8* expression in people with obesity was observed in both males and females (**Fig 1b**) and in individuals identifying as white, Black/African American, or Asian/Pacific Islanders (**Extended Data Fig 2**). Furthermore, *MXRA8* expression in WAT was positively correlated with whole-body adiposity (**Fig 1c**), subcutaneous abdominal white adipose tissue volume (**Fig 1d**), intra-abdominal adipose tissue volume (**Fig 1e**), and intrahepatic triglyceride content (**Fig 1f**). However, *MXRA8* expression in WAT did not correlate with lean body mass (**Fig 1g**) or bone mass (**Fig 1h**). These data indicate that MXRA8 expression is highly expressed in WAT and is increased in people with obesity, suggesting an important biological function for MXRA8 in adipose tissues.

**Figure 1.**
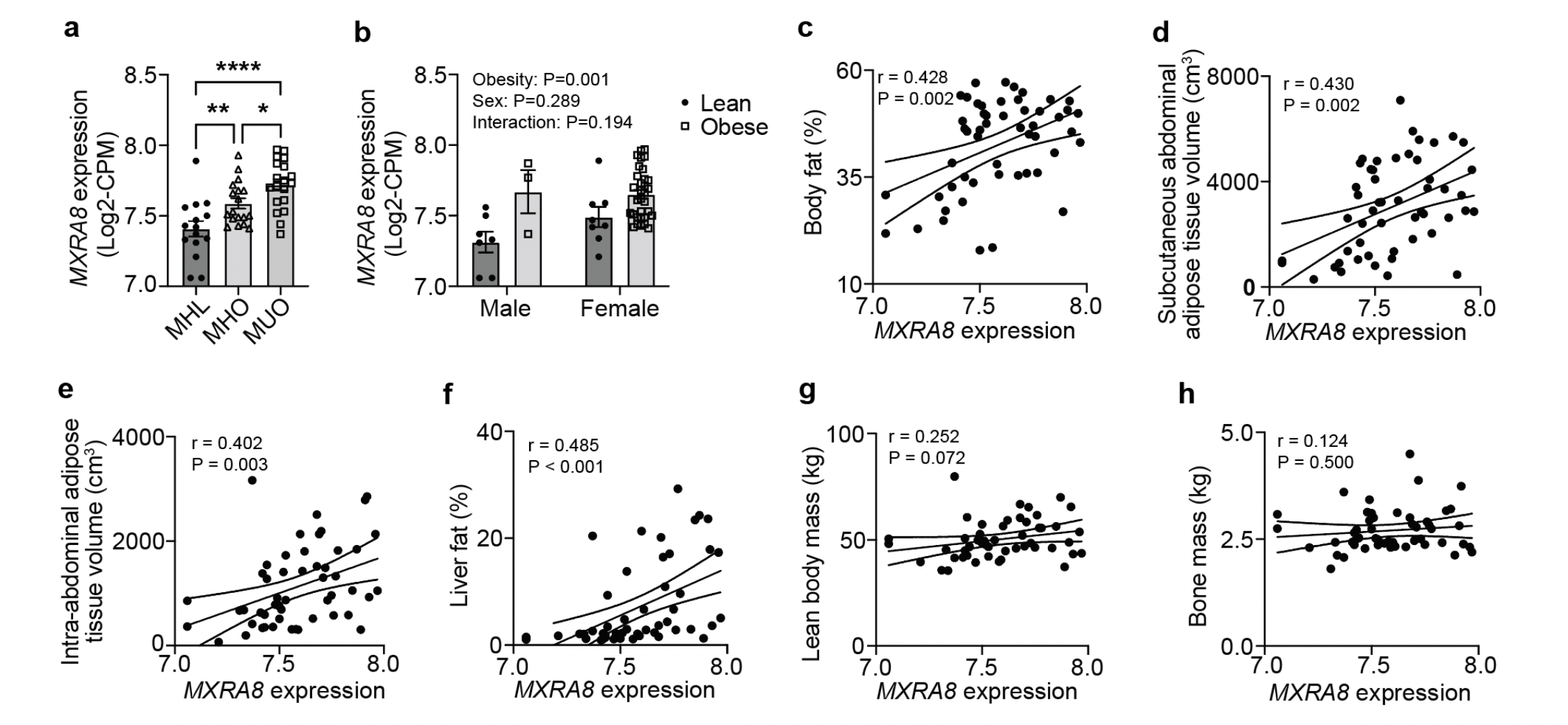
MXRA8 expression in white adipose tissue is upregulated in people with obesity and is associated with increased adiposity. Subcutaneous white adipose tissues (WAT) were obtained from people who were metabolically healthy lean (MHL, n=15), metabolically healthy obese (MHO, n=18), and metabolically unhealthy obese (MUO, n=19) for RNA-sequencing. **(a)** WAT *MXRA8* gene expression in MHL, MHO, and MUO groups. **(b)** WAT *MXRA8* expression in people who were lean or obese stratified by male or female biological sex, where the people with obesity included both MHO and MUO groups. **(c-h)** Linear regression analyses of WAT *MXRA8* expression and **(c)** adiposity expressed as percent body fat, **(d)** subcutaneous abdominal adipose tissue volume, **(e)** intra-abdominal adipose tissue volume, **(f)** liver fat, **(g)** lean body mass, and **(h)** bone mass. Data are expressed as mean ± standard error of the mean in panels a and b. For panel a, one-way ANOVA with Fisher’s LSD post hoc test. For panel b, two-way ANOVA with Fisher’s LSD post hoc test. For panels c-h, linear regression with Pearson correlation coefficients shown. *P<0.05, **P<0.01, ****P<0.0001.

As was the case in humans, mice also highly express MXRA8 in adipose tissues (**Fig 2a**). Epididymal (e)WAT and inguinal (i)WAT had significantly higher *Mxra8* transcript levels than brown adipose tissue (BAT), and all three adipose depots had substantially higher levels of *Mxra8* than did the spleen (**Fig 2a**), an organ that we expected to have moderate expression of MXRA8 based on the Human Protein Atlas dataset (ranked 10^th^ of 51 tissues; Extended Data Fig 1a). In addition, wildtype C57BL6/J (WT) mice fed a high fat diet (HFD) for 8 weeks had increased levels of MXRA8 protein in eWAT, iWAT, and BAT compared to control mice fed a normal chow diet (NCD, **Fig 2b-2c**). This result indicates that upregulation of MXRA8 expression in WAT is a conserved characteristic of obesity in mice and humans.

**Figure 2.**
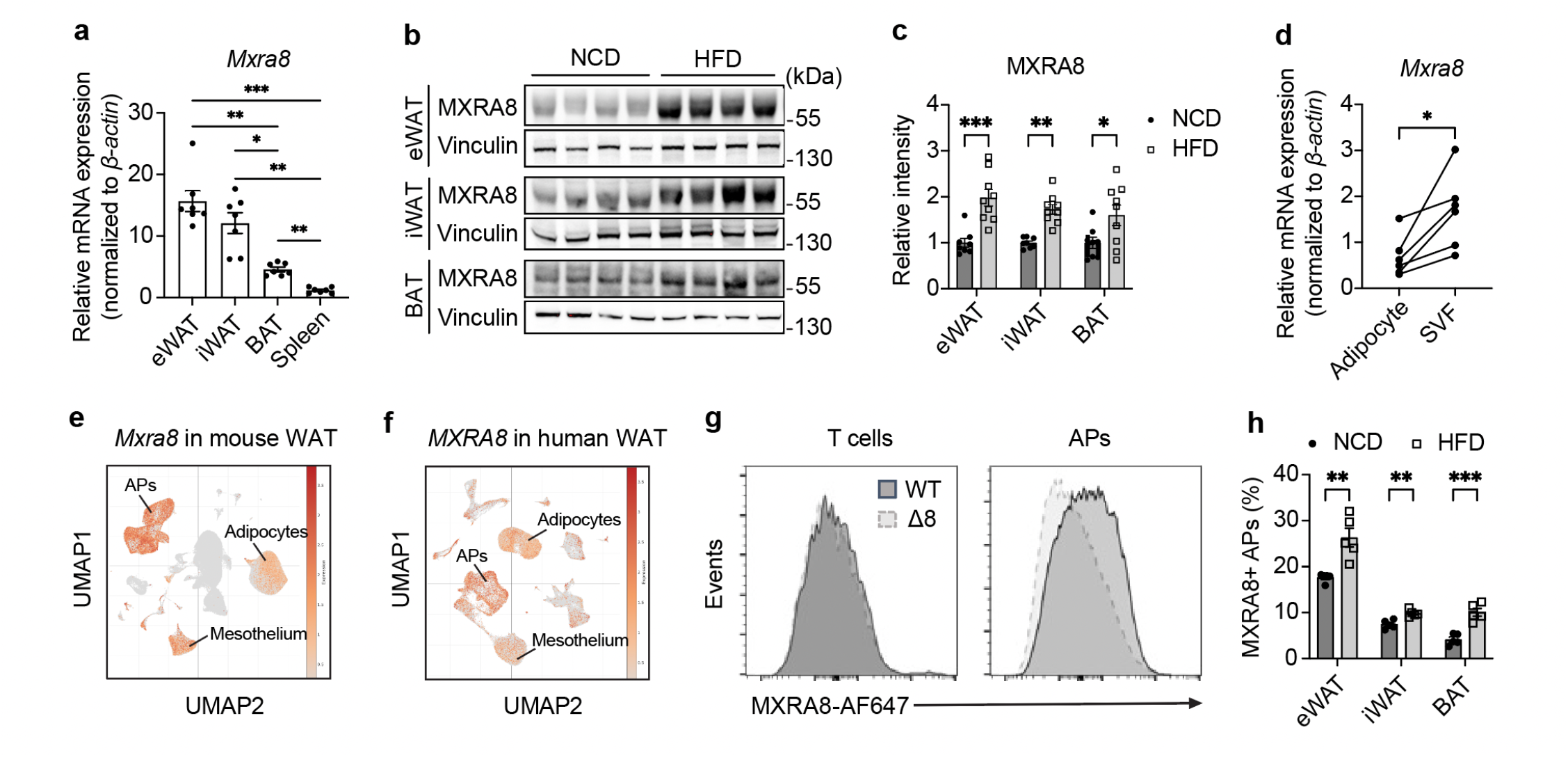
MXRA8 is highly expressed in adipocyte progenitor cells and is induced in obesity in mice. **(a)** Relative *Mxra8* mRNA expression in epididymal (e)WAT, inguinal (i)WAT, brown adipose tissue (BAT), and spleen from wildtype (WT, n=7) mice at age of 8-12-weeks-old. **(b)** Representative Western blot images and **(c)** densitometric quantification of MXRA8 protein levels in eWAT, iWAT, and BAT of WT mice that were fed a normal chow diet (NCD, n=8) or high fat diet (HFD, 60% kcal fat, n=9) for 8 weeks. Densitometric analyses are normalized to vinculin. **(d)** Relative *Mxra8* mRNA expression in mature, floating adipocytes and stromal vascular fraction (SVF) isolated from NCD-fed WT mice (n=6) at age of 8-12 weeks. **(e-f)** Uniform Manifold Approximation and Projection (UMAP) analyses on MXRA8 expression at a single cell level in **(e)** mouse and **(f)** human WAT based on a published single nucleus RNA sequencing dataset (GSE176171). **(g)** Representative flow cytometry histograms showing the surface expression of MXRA8 in T cells (gated as singlet live CD45^+^ NK1.1^−^ TCRαβ^+^) and adipocyte progenitor cells (APs, gated as singlet live CD45^−^ CD11b^−^ PDGFR1α^+^) in NCD-fed WT and Δ8 mice. **(h)** Frequencies of MXRA8 positive APs in eWAT, iWAT, and BAT of WT mice fed a NCD (n=5) or HFD (60% kcal fat, n=5) for 10 weeks starting at age 8-12-weeks-old. Data are expressed as mean ± standard error of the mean for panels a, c, and h. For panel d, paired specimens are linked with a line. For panel a, one-way ANOVA with Tukey post hoc test. For panel d, paired Student’s t-test. For panels c and h, two-way ANOVA with Sidak post hoc tests. *P<0.05, **P<0.01, ***P<0.001.

To determine the cell types that express MXRA8, we first isolated eWAT adipocytes and stromal vascular fraction (SVF) cells from WT mice and observed significantly higher expression of *Mxra8* transcripts in the SVF compared to the floating adipocytes (**Fig 2d**). As the SVF contains numerous cell types, including all immune cell lineages and adipocyte progenitor (AP) cells, we used publicly available single nucleus RNA sequencing datasets (GEO accession number GSE176171) and found that expression of MXRA8 mRNA was highest in AP cells in both mice (**Fig 2e**) and humans (**Fig 2f**). *MXRA8* mRNA was also expressed by mature adipocytes and mesothelial cells to lesser degrees in both species (**Fig 2e-2f**). To verify the presence of MXRA8 protein on the surface of AP cells, we conjugated anti-MXRA8 antibodies to Alexa Fluor 647 (AF647) and performed flow cytometry of eWAT SVF from WT vs *Mxra8*^*Δ8/Δ8*^ mice, which have an 8bp frameshift deletion after domain 2 of the *Mxra8* gene.^14^ We found that live CD45^−^ PDGFR1α^+^ AP cells were MXRA8^+^ in WT but not *Mxra8*^*Δ8/Δ8*^ mice (**Fig 2g** and **Extended Data Fig 3a**). Although some T cell subsets express MXRA8 in humans^5^, we did not observe MXRA8 expression on the surface of live CD45^+^ NK1.1^−^ TCRαβ^+^ T cells in eWAT from lean mice (**Fig 2g** and **Extended Data Fig 3a**). Next, we fed WT mice a normal chow diet (NCD) or HFD for 10 weeks and observed higher percentages of MXRA8^+^ AP cells in eWAT, iWAT, and BAT in the setting of HFD-induced obesity (**Fig 2h** and **Extended Data Fig 3b**). This result suggests that a subset of AP cells in white (eWAT), beige (iWAT), and brown fat (BAT) exhibit marked induction of MXRA8 expression in obese mice.

To determine whether MXRA8 has a functional role in adipose tissues, we compared the metabolic characteristics of MXRA8 mutant mice and WT littermate controls. First, we confirmed their genotypes in eWAT, iWAT, and BAT using primers that fail to amplify *Mxra8* mRNA if the 8bp deletion found in *Mxra8*^*Δ8/Δ8*^ mice is present (**Extended Data Fig 4**). On a normal chow diet, WT and *Mxra8*^*Δ8/Δ8*^ mice had similar body weights, lean mass, fat mass, and adiposity (**Extended Data Fig 5a-c**), and metabolic cage analyses indicated the two genotypes had similar energy expenditure, respiratory exchange ratios, activity levels, and food intake (**Extended Data Fig 5d-5h**). However, when fed a HFD *Mxra8*^*Δ8/Δ8*^ mice gained less weight (**Fig 3a**), accumulated less whole-body fat mass (**Fig 3b**), and had lower adiposity (**Fig 3c**) than WT controls. Although eWAT mass did not differ between groups, iWAT and BAT masses were significantly lower in *Mxra8*^*Δ8/Δ8*^ mice (**Fig 3d**). The reduction in fat mass accumulation was not limited to adipose tissues, as liver mass tended to be lower (**Fig 3e**) and liver adiposity was decreased in *Mxra8*^*Δ8/Δ8*^ mice (**Fig 3f**).

**Figure 3.**
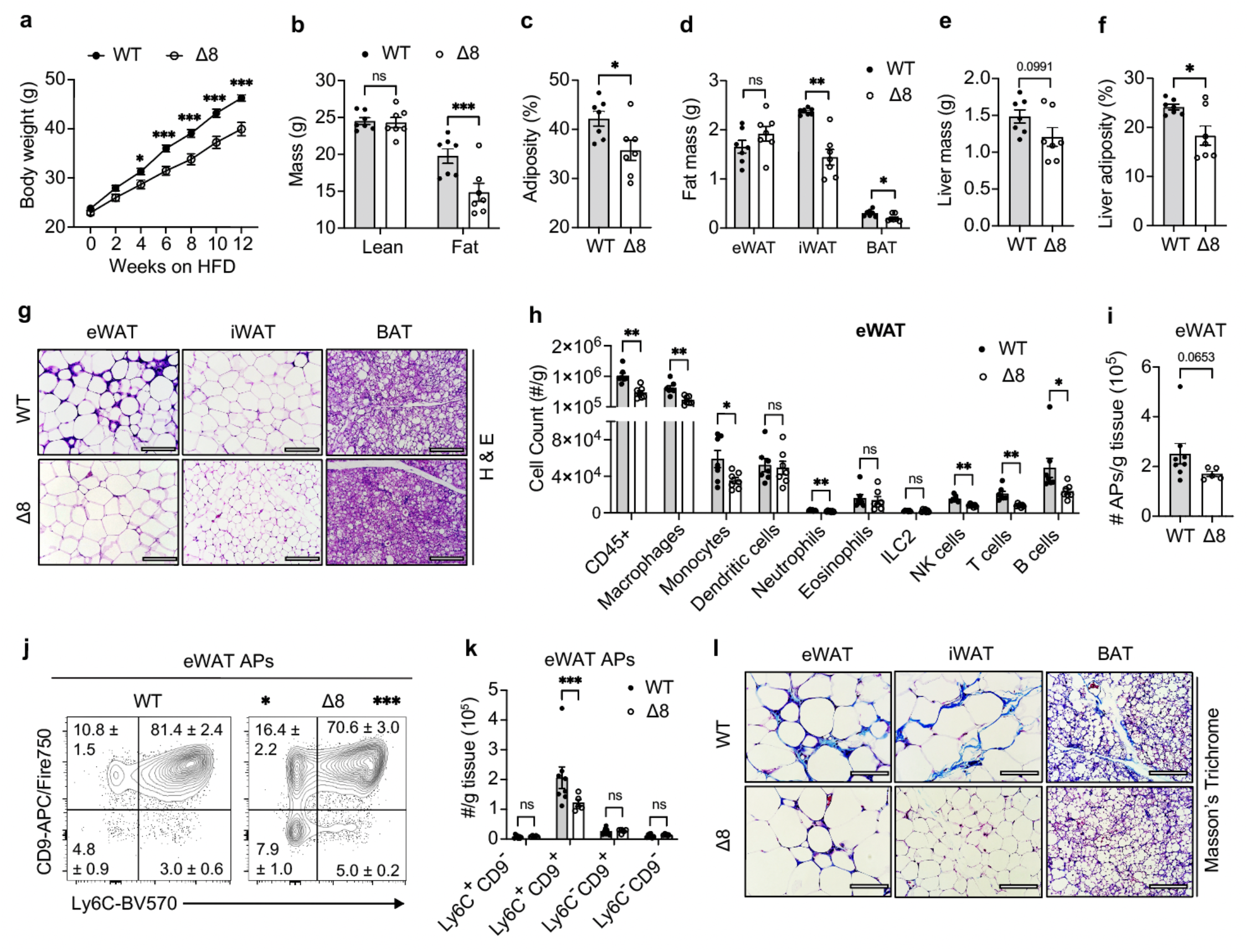
MXRA8 deficiency is associated with protection from HFD-induced obesity in mice. 8-10-week-old male wildtype (WT, n=7) or *Mxra8*^*Δ8/Δ8*^ (Δ8, n=7) mice were fed a HFD (60% kcal fat) for 12 weeks. **(a)** Body weight over time, **(b)** body composition analyses, **(c)** whole-body adiposity, **(d)** eWAT, iWAT, and BAT masses, **(e)** liver mass, and **(f)** liver adiposity. **(g)** Representative histologic images of eWAT, iWAT, and BAT with hematoxylin and eosin (H&E) staining, imaged at 400X magnification. Scale bar is 100 μm. **(h)** Numbers of total immune cells (CD45^+^) and immune cell populations per gram of eWAT based on spectral flow cytometric analyses. **(i-k)** 8-10-week-old male WT (n=8) and Δ8 (n=5) mice were fed a HFD for 12 weeks. **(i)** Numbers of singlet live CD45^−^ CD11b^−^ PDGFR1α+ adipocyte progenitor cells (APs) per gram of eWAT and **(j)** representative flow cytometry plots of eWAT AP cell subsets gated based on expression of CD9 and Ly6C. **(k)** Number of AP subsets per gram of eWAT. **(l)** Representative histologic images of eWAT, iWAT, and BAT with Masson’s Trichrome staining, imaged at 400X magnification. Scale bar is 100 μm. Data are expressed as mean ± standard error of the mean. For panels a, c, e, g, and j, two-way ANOVA with Fisher’s LSD post hoc test. For panel b, two-way ANOVA with repeated measures and Fisher’s LSD post hoc test. For panel d, Student’s t test. For panel h, Mann-Whitney U test. NS, not significant. *P<0.05, **P<0.01, ***P<0.001.

The reduction in iWAT and BAT mass but not eWAT mass in *Mxra8*^*Δ8/Δ8*^ mice fed a HFD suggested that MXRA8 may have adipose tissue-specific effects. Consistent with this possibility, histological analyses of eWAT, iWAT, and BAT revealed several prominent tissue-specific morphological features. In eWAT, there were no apparent differences in adipocyte size between groups, which is consistent with eWAT masses being similar. However, the eWAT contained numerous crown-like structures (CLS) in WT but not *Mxra8*^*Δ8/Δ8*^ mice fed a HFD (**Fig 3g**). CLS are highly enriched in immune cells and are strongly correlated with WAT inflammation in obesity.^15,16^ High-dimensional spectral flow cytometric analyses confirmed that eWAT from *Mxra8*^*Δ8/Δ8*^ mice had fewer total immune cells, macrophages, monocytes, neutrophils, natural killer/group 1 innate lymphoid cells, T cells and B cells compared to WT controls, whereas eosinophil and group 2 innate lymphoid cell abundances did not differ (**Fig 3h** and **Extended Data Fig 3a**). In iWAT, there were relatively few CLS in both genotypes, and the adipocytes were smaller with more abundant cytoplasm in *Mxra8*^*Δ8/Δ8*^ than WT mice (**Fig 3g**). These morphological features are suggestive of increased beige adipocytes in iWAT of *Mxra8*^*Δ8/Δ8*^ mice. Similarly, the BAT of *Mxra8*^*Δ8/Δ8*^ mice exhibited less lipid accumulation, smaller brown adipocytes, and more abundant cytoplasm compared to WT mice (**Fig 3g**).

The reduction in numerous immune cell populations in eWAT and high expression of MXRA8 on AP cells led us to examine whether MXRA8 regulates the abundance of an AP subset known as fibroinflammatory progenitors (FIPs). Recent studies indicate that FIPs are CD45^−^ PDGFR1α^+^ APs that express Ly6C and CD9 and produce numerous factors that drive inflammation and fibrotic remodeling of WAT in obesity.^17,18^ This tissue remodeling process is believed to be pathological and contribute to metabolic abnormalities in obese mice and humans, including impaired glucose homeostasis.^19,20^ There were fewer AP cells per gram of eWAT in *Mxra8*^*Δ8/Δ8*^ than WT mice fed a HFD (**Fig 3i**), and this change was explained by a decrease in the proportion and numbers (per gram of fat) of APs that were Ly6C^+^ CD9^+^ FIPs (**Fig 3j-3k**). Consistent with this result, there was reduced fibrosis in eWAT, iWAT, and BAT, as indicated by Masson’s trichrome staining in *Mxra8*^*Δ8/Δ8*^ mice compared to WT controls (**Fig 3l**). These data suggest that MXRA8 might regulate AP cell responses in adipose tissue and promote the inflammation and fibrosis that occur during tissue remodeling in obese WAT. However, it is also possible that these observed tissue remodeling phenotypes are explained by the reduction in weight gain in *Mxra8*^*Δ8/Δ8*^ mice. Further research is needed to understand how MXRA8 regulates the function of AP cells and associated tissue remodeling.

Next, we sought to understand the mechanisms by which *Mxra8*^*Δ8/Δ8*^ mice are protected from HFD-induced obesity. To begin studying this, we housed the mice in metabolic cages to characterize their energy expenditure and intake. We observed that *Mxra8*^*Δ8/Δ8*^ mice had significantly higher energy expenditure in both phases of the light:dark cycle (**Fig 4a-4b**). There were no differences in the respiratory exchange ratio, suggesting that energy substrate utilization was similar between the groups (**Fig 4c**). In addition, there were no significant alterations in activity levels (**Fig 4d**) or food intake (**Fig 4e**). These data suggest that *Mxra8*^*Δ8/Δ8*^ mice may have increased adaptive thermogenesis in setting of HFD feeding and that this process might contribute to their protection from diet-induced obesity.

**Figure 4.**
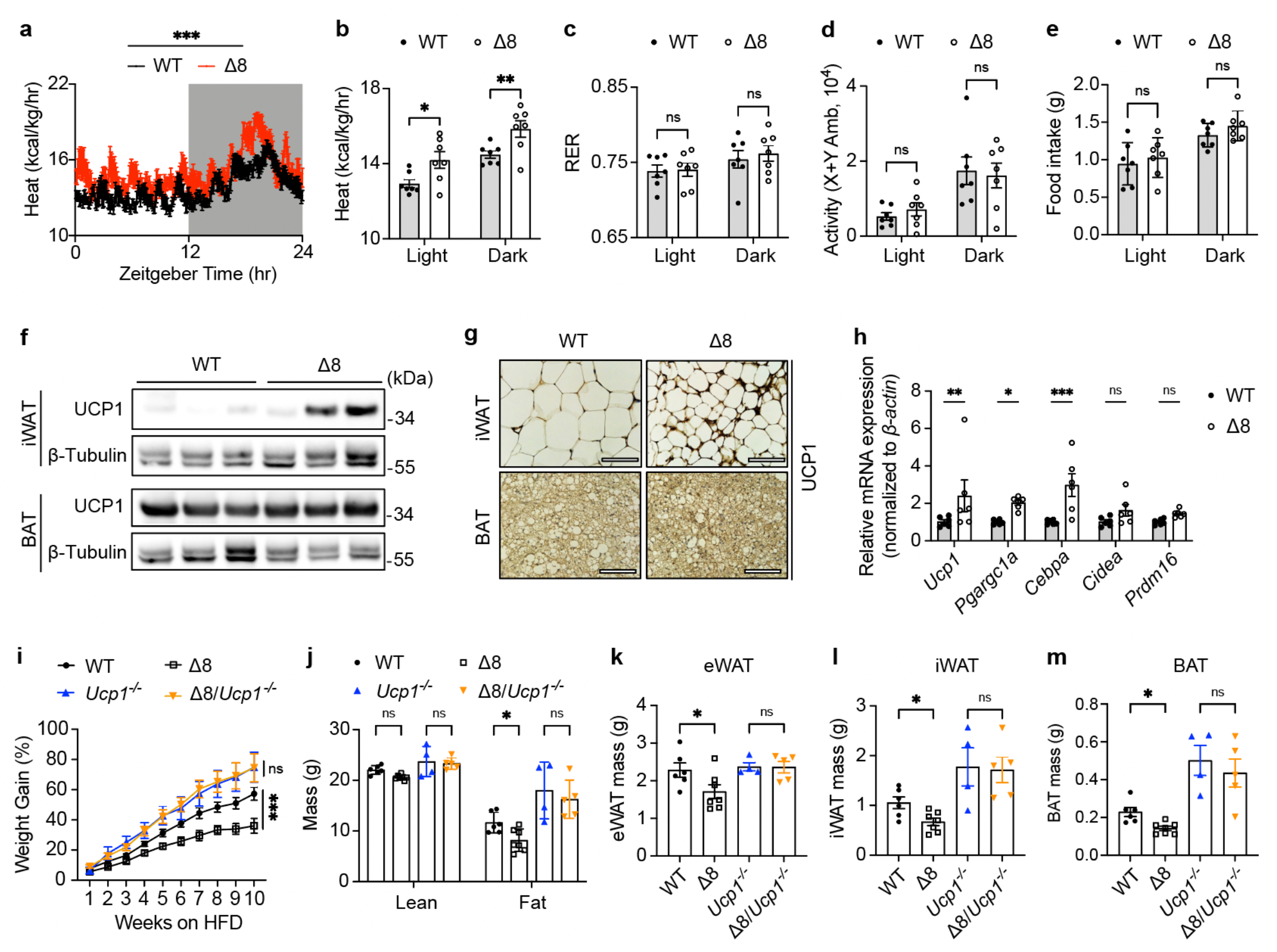
UCP1 is required for MXRA8 deficiency to protect against diet-induced obesity. **(a-g)** 8-10-week-old male wildtype (WT, n=7) and *Mxra8*^*Δ8/Δ8*^ (Δ8, n=7) mice were fed a high fat diet (HFD) for 12 weeks and housed in metabolic cages. **(a)** Energy expenditure over time, with the light phase unshaded and the dark phase shaded. **(b)** Average energy expenditure, **(c)** respiratory exchange ratio (RER), **(d)** ambulatory activity, and **(e)** food intake during the light and dark phases. **(f)** Representative Western blots of Uncoupling protein 1 (UCP1) in iWAT and BAT and **(g)** immunohistochemistry of UCP1 in iWAT and BAT. **(h)** Primary adipocytes were cultured from WT (n=6) or Δ8 (n=6) mice, and relative mRNA expression of the indicated genes on day 8. **(i-m)** 8-10-week-old male WT (n=6), Δ8 (n=7), *Ucp1*^*−/−*^ (n=4), and Δ8/ *Ucp1*^−/−^ (n=5) mice were acclimated to thermoneutrality (30°C) for 2 weeks prior to being fed a HFD (60% kcal fat) for 10 weeks at thermoneutrality. **(i)** Weight gain over time, expressed as a percentage of starting body weight. **(j)** Body composition analyses, **(k)** eWAT mass, **(l)** iWAT mass, and **(m)** BAT mass. Data are expressed as mean ± standard error of the mean. For panels a-e and i, two-way ANOVA with repeated measures and LSD post hoc tests. For panel h and j-m, two-way ANOVA with LSD post hoc test. NS, not significant. *P<0.05, **P<0.01, ***P<0.001.

One of the primary mechanisms of adaptive thermogenesis is UCP1-mediated heat generation. This protein is expressed by beige adipocytes in iWAT and brown adipocytes in BAT and is localized to the inner mitochondrial membrane, where UCP1 dissipates the proton gradient to generate large amounts of heat.^21^ UCP1^+^ beige and brown adipocytes thereby increase energy expenditure and limit the development of obesity in mice.^22,23^ However, beige and brown adipocytes undergo “whitening” in obese mice and humans, a process in which these cells lose their thermogenic capacity and downregulate expression of UCP1. Indeed, beige and brown adipocyte activation is severely impaired in obese mice and humans^21,24^, and it has been reported that the loss of beige and brown adipocytes is associated with an increased risk of developing cardiometabolic diseases in humans.^25^

As noted above, histological analyses of iWAT and BAT with hematoxylin and eosin staining suggested that *Mxra8*^*Δ8/Δ8*^ mice exhibited less whitening of iWAT (beige fat) and BAT (brown fat) (Fig 3). Consistent with this observation, UCP1 expression tended to be higher in iWAT and BAT of HFD-fed *Mxra8*^*Δ8/Δ8*^ mice than WT controls (**Fig 4f**). Furthermore, UCP1^+^ beige adipocytes were more abundant in iWAT, and there was increased UCP1^+^ signal intensity in brown adipocytes in BAT of *Mxra8*^*Δ8/Δ8*^ mice (**Fig 4g**). Although these results could be explained by the less severe obesity phenotype in *Mxra8*^*Δ8/Δ8*^ mice, an alternative possibility is that MXRA8 regulates the differentiation of thermogenic adipocytes. To investigate this question, we isolated AP cells from iWAT of WT or *Mxra8*^*Δ8/Δ8*^ mice and differentiated them into primary adipocytes. Indeed, we found that primary adipocytes from *Mxra8*^*Δ8/Δ8*^ mice exhibited significantly increased expression of *Ucp1* compared to WT adipocytes (**Fig 4h**). The transcription of *Ucp1* is mediated in part by the CCAAT/enhancer-binding protein α (C/EBPα, *Cebpa*)^26,27^ and peroxisome proliferator-activated receptor-γ coactivator (PGC)-1α (*Ppargc1a*)^28,29^, both of which were also increased in *Mxra8*^*Δ8/Δ8*^ primary adipocytes. The other thermogenic adipocyte-associated genes *Cidea* and *Prdm16* were not differentially expressed between groups. These transcriptional data suggest that MXRA8 inhibits the differentiation of AP cells into thermogenic adipocytes or regulates the expression of *Ucp1* in differentiated adipocytes.

To test whether UCP1-dependent thermogenesis is required for the weight-gain protection in MXRA8 mutant mice, we generated WT, *Ucp1*^*−/−*^, *Mxra8*^*Δ8/Δ8*^, and *Ucp1*^*−/−*^*;Mxra8*^*Δ8/Δ8*^ mice. We housed the 4 strains at thermoneutrality for 2 weeks prior to initiating a HFD because mice lacking UCP1 are profoundly cold-sensitive and activate behavioral and biochemical thermogenic mechanisms at room temperature that dramatically affect their response to diet-induced obesity.^30^ As had been reported previously^30-32^, we found that *Ucp1*^*−/−*^ mice gained more weight than WT controls when fed a HFD at thermoneutrality (**Fig 4i**). In addition, *Mxra8*^*Δ8/Δ8*^ mice were still protected from weight gain at thermoneutrality on a UCP1-sufficient background (**Fig 4i**). However, there were no differences in weight gain between the *Ucp1*^*−/−*^ and *Ucp1*^*−/−*^*;Mxra8*^*Δ8/Δ8*^ groups (**Fig 4i**). Body composition analyses further indicated that *Mxra8*^*Δ8/Δ8*^ mice accumulated less whole-body fat than WT mice, but this effect was ablated in the absence of UCP1 (**Fig 4j**). Similar trends were observed for eWAT, iWAT and BAT masses (**Fig 4k-4m**). Collectively, these data indicate that the MXRA8 mutant mice are protected against HFD-induced weight gain in a UCP1-dependent manner.

In summary, we find that MXRA8 is highly expressed in WAT and upregulated in this tissue in obesity in both mice and humans. This protein is primarily expressed by AP cells and adipocytes and functions, in part, to inhibit the differentiation or thermogenic potential of beige and/or brown adipocytes. Mice that lack intact, full-length MXRA8 (*Mxra8*^*Δ8/Δ8*^) are protected from HFD-induced obesity, exhibit decreased iWAT (beige fat) and BAT (brown fat) masses, have reduced fibroinflammatory changes that are characteristic of pathological adipose tissue remodeling in obesity, and have increased UCP1 expression in iWAT and BAT. Furthermore, expression of UCP1 was required for *Mxra8*^*Δ8/Δ8*^ mice to be protected from HFD-induced obesity.

These findings suggest a model in which upregulation of MXRA8 promotes whitening of beige and brown adipose tissues in obesity, leading to a loss of UCP1-mediated thermogenic potential of adipose tissues and increased obesity in mice. Although our data suggest that MXRA8 impairs the differentiation or thermogenic function of adipocytes, there may be additional mechanisms by which MXRA8 regulates adipose tissue physiology and obesity pathogenesis. One possibility is that MXRA8 may lead to whitening of beige and brown adipose tissues by disrupting angiogenesis in adipose tissues. Indeed, MXRA8 has been reported to inhibit angiogenesis^6^, and the loss of vascular density drives whitening in beige and brown adipose tissues in obesity.^33,34^ Another possibility is that MXRA8 may bind extracellular matrix proteins or integrins, as was suggested for α_V_β_3_ integrin^2^, to mediate adipose tissue remodeling and the progression of metabolic dysfunction in obesity. This possibility is supported by our observation that WAT from *Mxra8*^*Δ8/Δ8*^ mice had markedly reduced fibrosis compared to WT controls. Additional studies are needed to determine the role of MXRA8 in regulating angiogenesis and remodeling of the extracellular matrix in adipose tissues.

MXRA8 has been implicated in several diseases, including the progression of several types of cancer^35,36^, colitis^37^, and autoimmunity^5^, and also serves as an entry receptor for some arthritogenic alphaviruses, such as chikungunya virus.^7^ Our studies are the first to demonstrate a role for MXRA8 in regulating adipose tissue function and obesity pathogenesis and suggest that MXRA8 may be a previously unappreciated therapeutic target to treat obesity and potentially other metabolic diseases.

## Acknowledgements

J.R.B. is supported by the National Institutes of Health (NIH) Office of the Director (DP5 OD028125) and Burroughs Wellcome Fund (CAMS #1019648). W.J. is supported by an American Heart Association (AHA) Postdoctoral Fellowship (24POST1244220). R.G. is supported by an AHA Predoctoral Fellowship (24PRE1189775). J.M.W. is supported by NIH T32 AI007163. S.K. is supported by a W.M. Keck Foundation Postdoctoral Fellowship in Molecular Medicine. The study was supported by the Musculoskeletal Research Center histology core facility (P30 AR074992) and the Nutrition Obesity Research Center (P30 DK56341).

## Author Contributions

WJ performed experiments, analyzed data, interpreted results, and wrote the manuscript. RG, JRM, OA, RLF, SV, SRC, JW, XZ, and SK performed experiments, analyzed data, and interpreted results. GIS, MP, and SK performed experiments, analyzed data, and interpreted results. IJL and NAA contributed to experimental designs and interpreted results. MSD and ASK generated and provided mouse strains and reagents, provided guidance on experimental design, and interpreted results. JRB conceived of the project, secured funding, performed experiments, interpreted results, and wrote the manuscript. All authors contributed to writing, editing, and/or revising this manuscript and approved the final version.

## Data Availability Statement

All data supporting the findings of this study are available within the paper and its Supplementary Information or are available by request to the corresponding author.

## METHODS

### Human subjects

A total of 52 males and females participated in this study. The following criteria were used for inclusion in one of the three groups: i) metabolically healthy lean (MHL, n=15) defined as having a body mass index (BMI) of 18.5-24.9 kg/m^2^, and normal fasting plasma glucose (<100 mg/dL), oral glucose tolerance (2-h glucose <140 mg/dL), liver fat content (<5%), plasma triglycerides (<150 mg/dl), and normal whole-body insulin sensitivity, defined as the glucose infusion rate (GIR) per kg fat-free mass divided by the plasma insulin concentration (GIR/I) during a hyperinsulinemic-euglycemic clamp procedure (>40 μg/kg FFM/min/μU/mL); metabolically healthy obese (MHO, n=18) defined as having a BMI of 30.0-49.9 kg/m^2^ and normal fasting plasma glucose, oral glucose tolerance, plasma triglycerides, and liver fat content and normal whole-body insulin sensitivity; and metabolically unhealthy obese (MUO, n=19) defined as having a BMI of 30.0-49.9 kg/m^2^, impaired fasting glucose or oral glucose tolerance, high liver fat content (≥6%) and impaired whole-body insulin sensitivity, defined as a GIR/I ≤40 (μg/kg FFM/min)/(μU/mL). All subjects provided written, informed consent before participating in this study, which was approved by the Institutional Review Board of Washington University School of Medicine in St. Louis, MO and registered in ClinicalTrials.gov (NCT02706262). The assessments of body composition, oral glucose tolerance, and insulin sensitivity, and the acquisition of periumbilical subcutaneous abdominal adipose tissue for bulk RNA-sequencing analyses were conducted as previously described.^13^ *MXRA8* mRNA counts were extracted from the RNAseq dataset for targeted analyses.

### Mice

*Mxra8*^*Δ8/Δ8*^ (Δ8) mice were generated as previously described^14^, and were maintained by crossing *Mxra8*^*+/Δ8*^ to *Mxra8*^*+/Δ8*^ mice to generate wildtype (WT) and Δ8 littermates. *Ucp1*^−*/*−^ mice were obtained from Jackson Laboratories (strain #003124) and crossed Δ8 mice to generate *Mxra8*^*Δ8/Δ8*^*/Ucp1*^−*/*−^ (Δ8/*Ucp1*^−*/*−^) mice. These double mutants were maintained by homozygous breeding, as were their single-gene knockout counterparts, with timed breeding to ensure that cohorts of cousins were the same age. Wildtype C57BL6/J mice were either purchased from The Jackson Laboratory (strain #000664) and bred in-house for experimental use. All mice were housed in a specific pathogen-free facility with a 12h:12h light:dark cycle (lights on from 06:00am to 06:00pm) and *ad libitum* access to food and water. Normal Chow Diet was utilized for routine feeding, and 60% kcal fat HFD made from lard (cat# D12492, Research Diets, Inc.) was employed for establishing diet-induced obese mouse models with the indicated feeding timelines. Studies involving *Ucp1*^−*/*−^ strains were singly housed at thermoneutrality (30°C) for 2 weeks prior to HFD feeding.^30,38^ All other mice were housed in groups at room temperature (22°C). Animals were randomly assigned to n=2-5 mice/group per experiment depending on the numbers of available mice, and data from at least 2 independent experiments were pooled for analyses. Mice were euthanized using isoflurane inhalation immediately prior to harvesting inguinal white adipose tissue (iWAT), epididymal (e)WAT (males), ovarian (o)WAT (females), interscapular brown adipose tissue (BAT), liver, and spleen. All experiments were performed according to the guidelines of the Institutional Animal Care and Use Committee (IACUC) at Washington University in St. Louis and in accordance with IACUC-approved protocol 22-0286.

### Metabolic cage analyses

Metabolic cage analyses were performed using a 16-metabolic cage Comprehensive Laboratory Animal Monitoring System (CLAMS) (Columbus Instruments, Columbus, OH) as previously described.^39^ Briefly, mice were weighed and body composition was measured using an EchoMRI 2n1 with a horizontal configuration. The mice were placed in the CLAMS cages (one mouse per cage) with a 12h:12h light-dark cycle and were provided free access to food and water, both of which were hung on a load cell. Mice were arranged in a staggered manner in CLAMS to ensure horizontally and vertically equal distribution of groups. Mice were allowed to acclimate for 1 day. Data were analyzed on the first full 24h period inclusive of a complete 12h light phase followed by a complete 12h dark phase. During the measurement period, cumulative food and water intake, spontaneous activity, oxygen (O_2_) consumption and carbon dioxide (CO_2_) production were monitored. The respiratory exchange ratio (RER) was calculated by dividing the volume of CO_2_ produced by the volume of O_2_ consumed. Energy expenditure was computed using the standard equation and normalized to body weight.

### Body composition and liver adiposity analyses in mice

Body composition was measured using an EchoMRI-100H 2n1 featuring a horizontal probe configuration (EchoMRI, Houston, TX). Whole-body adiposity was calculated as the ratio of fat mass to body weight. To determine liver adiposity, the entire liver was harvested, weighed, and analyzed using the EchoMRI 2n1 tissue probe. Liver fat mass was divided by liver mass to calculate liver adiposity.

### Isolation of stromal vascular fraction from adipose tissues

The stromal vascular fraction (SVF) from mouse adipose tissues were isolated as previously described.^39,40^ After euthanasia, eWAT, iWAT, and BAT were immediately dissected, finely minced, and subjected to digestion in 4 mL high glucose DMEM (Gibco) containing 1 mg/mL collagenase type II (cat# C6885, Sigma-Aldrich). Digestion was carried out at 37 °C for 1 hr in an orbital shaker with rotation at 140 rpm while tilted at a 90° angle. The resulting single-cell suspensions were filtered through a 100 μm nylon mesh cells strainer followed by two washes of filter with 5 mL of Wash Media (high glucose DMEM with 5% FBS, 2 mM L-glutamine, and 100 U/mL Penicillin-Streptomycin). After centrifugation at 500 x *g* for 5 min at 4°C, the floating adipocytes on the top of the media were removed (in some cases they were collected), as was the media. The SVF pellet was resuspended in 1 mL of Red Blood Cell ACK Lysis Buffer (Gibco) and incubated at room temperature for 3-5 min to lyse red blood cells. After quenching with 10 mL Wash Media, SVF cells were pelleted by centrifugation at 500 x *g* for 5 min at 4°C and resuspended in appropriate volume of wash media for subsequent experiments such as flow cytometry staining and primary adipocyte differentiation.

### Spectral flow cytometry

Cell staining for spectral flow cytometry was performed as previously described.^40^ Isolated SVF cells were plated into 96-well round-bottom plates, washed with 200 μL DPBS, and then resuspended in 50 μL Zombie Near Infrared (Zombie-NIR; 1:1,000; BioLegend) in DPBS. After a 5 min incubation on ice while protected from light, the Zombie viability dye was quenched with 200 μL FACS Buffer (DPBS supplemented with 2.5% heat-inactivated FBS and 2.5 mM EDTA). Cells were then pelleted at 500 x *g* for 3 min at 4°C and resuspended in 25 μL of 5 mg/mL FcBlock (rat anti-mouse CD16/32, clone 2.4G2, BD Biosciences) diluted in FACS Buffer and incubated on ice for 10-15 min. An equal volume of 2X stain cocktail was made in Brilliant Stain Buffer (BD Biosciences) and added on top, with gentle mixing by pipetting up and down 4-5 times. The 2X stain cocktail included the following antibodies (final dilutions are all 1:300 unless otherwise indicated): rat anti-mouse SiglecF-BV421 (1:400, clone E50-2440, cat# 562681 BD Horizon), rat anti-mouse/human CD11b-Pacific Blue (clone M1/70, cat# 101224, BioLegend), rat anti-mouse ST2/IL-33R-biotin (clone DIH9, cat# 145308, BioLegend), rat anti-mouse MHC-II-BV510 (clone M5/114.15.2, cat# 107636, BioLegend), rat anti-mouse Ly6C-BV570 (1:400, clone HK1.4, cat #128030, BioLegend), rat anti-mouse F4/80-BV650 (clone BM8, cat#123149, BioLegend), rat anti-mouse Ly6G-BV711 (clone 1A8, cat# 127643, BioLegend), Armenian hamster anti-mouse TCRβ-Alexa Fluor 488 (clone H57-597, cat# 109215, BioLegend), rat anti-mouse CD45-PerCP (1:200, clone 30-F11, cat#103130, BioLegend), mouse anti-mouse CD64-PE/Dazzle594 (clone X54-5/7.1, cat# 139320, BioLegend), rat anti-mouse PDGFRα-PE/Cy5 (clone APA5, cat# 135920, BioLegend), Armenian hamster anti-mouse CD11c-PE/Cy5.5 (clone N418, cat#35-0114-82, Invitrogen/eBioscience), rat anti-mouse CD25-PE/Cy7 (1:200, clone PC61, cat#1026, BioLegend), Armenian hamster anti-mouse MXRA8-Alexa Fluor 647 (1:200, clone 9G2.D6, conjugated using the Alexa Fluor™ 647 Antibody Labeling Kit by Invitrogen), rat anti-mouse CD19-Spark NIR 685 (clone 6D5, cat# 115568, BioLegend), rat anti-mouse CD9-APC/Fire 750 (clone MZ3, cat# 124814, BioLegend) or mouse anti-mouse NK1.1-APC/Fire750 (clone PK136, cat# 108752, BioLegend), and rat anti-mouse/human B220-APC/Fire810 (clone RA3-6B2, cat# 103278, BioLegend) in Brilliant Stain Buffer (BD Biosciences) supplemented with 5 μg/mL FcBlock. Cells were stained for 30 min on ice while protected from light followed by 2-to-3 washes in 200 μL FACS Buffer. Cells were then stained with 50 μL Streptavidin-BV480 (1:300, BioLegend) in FACS Buffer for 20 min on ice while protected from light. After 2 washes in 200 μL FACS Buffer, the cells were resuspended in 200 μL FACS Buffer and subjected to flow cytometric analysis on a Cytek Aurora spectral flow cytometer configured with 4 lasers (violet, blue, yellow/ green, and red lasers), with 100 μL acquired to enable cell count enumeration.

### Primary Adipocyte Differentiation and Culture

Freshly isolated SVF from iWAT of *Mxra8*^*Δ8/Δ8*^ and WT littermate controls were resuspended in preadipocyte culture media (DMEM/F12 (1:1) supplemented with 10% heat-inactivated FBS, 2 mM L-glutamine, and 100 U/mL Penicillin-Streptomycin) and cultured in a humidified incubator at 37°C with 5% CO_2_. After 2 days growth, cells were washed twice with sterile DPBS and lifted by 0.05% trypsin-0.44 mM EDTA in DPBS (Gibco) with 3-minute incubation at 37°C. The trypsinization was quenched by collecting the cells into a 50 mL conical tube containing 5 mL preadipocyte culture media. Cells were then pelleted by centrifugation at 400 x *g* for 3 min at 4°C, resuspended in preadipocyte culture media, and counted using a Countess II (Invitrogen). 3 × 10^5^ cells were plated into each well of a 6-well plate and maintained in a humidified incubator at 37°C with 5% CO_2_. Two days after the cell confluency reached 100%, the differentiation was initiated by replacing the culture media with adipocyte differentiation media (DMEM/F12 (1:1) supplemented with 10% heat-inactivated FBS, 5 μM dexamethasone (Sigma), 850 nM insulin (Sigma), 1 μM rosiglitazone (Cayman Chemical), 1 nM 3,3′,5-Triiodo-L-thyronine (Sigma), 125 μM indomethacin (Sigma), 0.5 mM 3-Isobutyl-1-methylxanthine (Sigma), and 100 U/mL Penicillin-Streptomycin) for 48 hr. The adipocyte differentiation media was then replaced by maintenance media (DMEM/F12 (1:1) supplemented with 10% heat-inactivated FBS, 850 nM insulin, 1 nM 3,3′,5-Triiodo-L-thyronine, and 100 U/mL Penicillin-Streptomycin). Primary adipocytes were obtained after 8 days incubation in maintenance media (refreshed every 2 days) and were harvested for RNA extraction and RT-qPCR using TRIzol.

### Histological Analyses

Freshly dissected eWAT, iWAT, BAT, and liver tissues were immediately immersed in 4% paraformaldehyde (PFA) in PBS (cat# sc-281692, Santa Cruz Biotechnology, Dallas, TX) and kept at 4°C for at least 3 days while protected from light. Fixed tissues were transferred into 75% ethanol and submitted to the WashU Musculoskeletal Research Center Morphology Core for paraffin embedding, sectioning, and hematoxylin and eosin (H&E) staining and Masson’s Trichrome staining. Unstained sections were used for immunohistochemistry (IHC) staining with the use of ImmPRESS HRP Universal PLUS Polymer Kit (cat# MP-7800, Vector Laboratories,). Briefly, the slides were sequentially subjected to dehydration in xylene and rehydration in 100% ethanol, 95% ethanol, 75% ethanol, and distilled water followed by heat-induced antigen retrieval in citrate buffer (cat# C9999, Sigma-Aldrich) using a 2100 Retriever (Electron Microscopy Sciences). After citrate buffer cooled down, slides were rinsed 3 times in Tris-Buffered Saline (TBS) and then incubated in BLOXALL Endogenous Enzyme Blocking Solution (cat# SP-6000, Vector Laboratories) for 15 min in dark to quench endogenous peroxidase. Slides were then rinsed 3 times in TBS and incubated in 2.5% Normal Horse Serum for 30 min at room temperature. The slides were stained overnight with rabbit anti-mouse UCP1 antibody (cat# ab10983, Abcam) diluted in 2.5% Normal Horse Serum (1:1000) at 4°C. Sections were rinsed 3 times in TBS and incubated in ImmPRESS HRP Universal Polymer Reagent (Horse Anti-Mouse/Rabbit IgG) for 30 minutes at room temperature. After 3 washes with TBS, ImmPACT DAB EqV working solution (mixture of equal volume of ImmPACT DAB EqV Reagents 1 and 2) were gently pipetted onto the sections and incubated until sharp brown signals developed, typically within 1-2 min. Slides were then rinsed in tap water 3 times and counterstained with hematoxylin (cat# H-3401, Vector Laboratories). Images were captured with a 40X Apochromat N.A. 0.95 objective using an Echo Rebel brightfield microscope (Discover ECHO, San Diego, CA) configured with a 10X flip-out Achromat condenser.

### RNA extraction and quantitative RT-PCR from mouse specimens

Total RNA was extracted from mouse tissues or cells, including mature adipocytes, SVF, and primary adipocytes, using the Direct-zol RNA Miniprep Plus Kit (Zymo Research; Orange, CA) based on the manufacturer’s instructions, with a chloroform extraction step added to remove the lipids prior to application to the columns. Frozen mouse tissues were lysed and homogenized in Direct-zol by a Bead mill homogenizer (Thermo Fisher, Waltham, MA). Cells were lysed by directly adding Direct-zol into the collecting tubes or culture wells. A half volume of chloroform was then added into the lysates, thoroughly mixed by vortexing, and centrifuged for 15 min at 16,000 x *g* at 4°C. The clear aqueous phase was collected into a fresh set of Eppendorf tubes that contained equal volumes of 100% molecular grade ethanol and gently mixed by inversion followed by applying the mixture onto the Zymo-Spin IIICG column for centrifugation. Total RNA was then purified according to the manufacturer’s instructions and quantified by absorbance on a BioTek Synergy H1 microplate reader (Biotek). Depending on the amount of RNA obtained, 100-2000 ng RNA were reverse-transcribed into cDNA using SuperScript™ IV VILO™ Master Mix (Applied Biosystems). Quantitative real-time PCR was performed using PowerUp™ SYBR™ Green Master Mix (Applied Biosystems) and on a QuantStudio 3 Real-Time PCR System (Applied Biosystems). The primers were all purchased from Millipore Sigma with sequences listed as below, *Mxra8(8bp)*-fwd: 5’-TCGTGCTTCTCCTGGCAATG-3’, *Mxra8(8bp)*-rev: 5’-GGAAGAAATGTGTGTGGTCCTC-3’; *Ucp1*-fwd: 5’-CAACTTGGAGGAAGAGATACTGAACAT-3’, *Ucp1*-rev: 5’-TTTGGTTGGTTTTATTCGTGGTC-3’; *Cebpa*-fwd: 5’-TTCACATTGCACAAGGCACT-3’, *Cebpa*-rev: 5’-GAGGGACCGGAGTTATGACA-3’; *Cidea*-fwd: 5’-TGCTCTTCTGTATCGCCCAGT-3’, *Ppargc1a*-fwd: 5’-CCCTGCCATTGTTAAGACC-3’, *Ppargc1a*-rev: 5’-TGCTGCTGTTCCTGTTTTC-3’; *Cidea*-rev: 5’-GCCGTGTTAAGGAATCTGCTG-3’; *Prdm16*-fwd: 5’-CAGCACGGTGAAGCCATTC-3’, *Prdm16*-rev: 5’-GCGTGCATCCGCTTGTG-3’; and *β-actin*-fwd: 5’-CTAAGGCCAACCGTGAAAAG-3’, *β-actin*-rev: 5’-ACCAGAGGCATACAGGGACA-3’.

### Protein Extraction and Western Blotting

Total proteins were extracted from eWAT, iWAT, and BAT using the Minute™ Total Protein Extraction Kit for Adipose Tissue/Cultured Adipocytes (Invent Biotechnologies; Plymouth, MN) according to the manufacturer’s instructions. Homogenization/lysis buffer was prepared prior to use by adding Halt™ Protease and Phosphatase Inhibitor Single-Use Cocktail (Thermo Scientific) and 1mM PMSF (Cell Signaling Technology; Danvers, MA) into the Buffer A provided by the above-mentioned kit. Proteins were precipitated from the lysates by acetone with overnight incubation at -20°C and resolubilized in 50-200 μL of resolubilization buffer (1% SDS, 1 mM EDTA, and 100 mM HEPES (pH7.5) in molecular biology-grade water). BCA Protein Assay Kit (Thermo Fisher; Waltham, MA) was utilized for protein quantification. For SDS-PAGE, 10 μg protein from each sample were prepared in Bolt™ LDS Sample Buffer (Invitrogen) under reducing (with 1 mM DTT) conditions and were heated at 70 °C for 10 min. Protein samples were then electrophoresed using Bolt™ 4-12% Bis-Tris Plus Gels (Invitrogen) and transferred onto nitrocellulose membranes using a Power Blotter System (Invitrogen). After washing with deionized water, membranes were stained with 0.01 % (w/v) Ponceau S (Sigma) in 1% aqueous acetic acid and imaged in an iBright CL1500 imaging system (Invitrogen). Membranes were then washed 3 times in Tris-buffered saline containing 0.1% Tween-20 (TBST, Cell Signaling Technology) prior to 1-hr blocking with blocking buffer (5% Blotting Grade Blocker (Bio Rad; Hercules, CA) in TBST) at room temperature. Membranes were probed overnight at 4°C with Armenian hamster anti-mouse MXRA8 (Clone 9G2.D6, 0.5 μg/ml), rabbit anti-mouse UCP1 (cat# ab10983, 1:1000, Abcam), β-tubulin (clone 9F3, cat# 2128, Cell Signaling Technology), or Vinculin (clone E1EV9, cat# 13901, Cell signaling Technology) diluted in blocking buffer supplemented with 0.01% NaN_3_ (Sigma). After three washes with TBST, membranes were incubated with horseradish peroxidase (HRP)-conjugated goat anti-rabbit (cat# 7074, Cell Signaling Technology) or goat anti-hamster (cat# 127-035-160, Jackson ImmunoResearch) secondary antibodies for 1 hr at room temperature. Blots were then developed using SuperSignal West Pico Chemiluminescent Substrate (Invitrogen) or SuperSignal West Atto Chemiluminescent Substrate (Invitrogen) and imaged with an iBright CL1500 Imaging System (Invitrogen). The densitometric analyses of Western blot bands was performed on iBright Analysis Software version 5.2.0. Vinculin or β-tubulin served as internal reference control for the normalization of target protein bands. Relative intensities were calculated by dividing each normalized intensity by the average normalized intensity of the control (WT) lanes.

### Statistical analyses

Data are presented as mean ± standard error of the mean in all panels with a pool from at least 2 independent experiments. Statistical analyses were performed in Prism v10 or v11 (Graphpad, La Jolla, CA) unless otherwise specified. Paired or unpaired Student’s t-tests were used for two-group comparisons, with Welch’s correction applied when the standard deviations between groups differed. One-way analysis of variance (ANOVA) with Tukey or Fisher’s LSD post-hoc testing was used for three-or-more group comparisons, and two-way ANOVA with Sidak or Fisher’s LSD posthoc testing was used for 2 × 2 or 2 x n designs, including experimental designs that involve repeated measures. Metabolic cage analyses were performed on OxyMax CI-Link software for CLAMS Cages. Immunoblot densitometric analyses were conducted on iBright CL1500 software. Spectral flow cytometry data were acquired using SpectroFlo v2.0 (Cytek) and analyzed on SpectroFlow v2.0 and FlowJo v10.8.1 (BD). Statistical significance was set at P<0.05.

## REFERENCES

1 Jung, Y. K. et al. DICAM, a novel dual immunoglobulin domain containing cell adhesion molecule interacts with alphavbeta3 integrin. J Cell Physiol 216, 603-614, doi:10.1002/jcp.21438 (2008).

2 Jung, Y. K. et al. DICAM inhibits osteoclast differentiation through attenuation of the integrin alphaVbeta3 pathway. J Bone Miner Res 27, 2024-2034, doi:10.1002/jbmr.1632 (2012).

3 Han, S. et al. Dicam promotes proliferation and maturation of chondrocyte through Indian hedgehog signaling in primary cilia. Osteoarthritis Cartilage 26, 945-953, doi:10.1016/j.joca.2018.04.008 (2018).

4 Simpson, K. E., Staikos, C. A., Watson, K. L. & Moorehead, R. A. Loss of MXRA8 Delays Mammary Tumor Development and Impairs Metastasis. Int J Mol Sci 24, doi:10.3390/ijms241813730 (2023).

5 Charabati, M. et al. DICAM promotes TH17 lymphocyte trafficking across the blood-brain barrier during autoimmune neuroinflammation. Sci Transl Med 14, eabj0473, doi:10.1126/scitranslmed.abj0473 (2022).

6 Han, S. W. et al. DICAM inhibits angiogenesis via suppression of AKT and p38 MAP kinase signalling. Cardiovasc Res 98, 73-82, doi:10.1093/cvr/cvt019 (2013).

7 Zhang, R. et al. Mxra8 is a receptor for multiple arthritogenic alphaviruses. Nature 557, 570-574, doi:10.1038/s41586-018-0121-3 (2018).

8 Basore, K. et al. Cryo-EM Structure of Chikungunya Virus in Complex with the Mxra8 Receptor. Cell 177, 1725–1737 e1716. doi:10.1016/j.cell.2019.04.006 (2019).

9 Song, H. et al. Molecular Basis of Arthritogenic Alphavirus Receptor MXRA8 Binding to Chikungunya Virus Envelope Protein. Cell 177, 1714-1724 e1712, doi:10.1016/j.cell.2019.04.008 (2019).

10 Cohen, P. et al. Ablation of PRDM16 and Beige Adipose Causes Metabolic Dysfunction and a Subcutaneous to Visceral Fat Switch. Cell 156, 304-316, doi:10.1016/j.cell.2013.12.021 (2014).

11 Wu, J. et al. Beige Adipocytes Are a Distinct Type of Thermogenic Fat Cell in Mouse and Human. Cell 150, 366-376, doi:10.1016/j.cell.2012.05.016 (2012).

12 Sidossis, L. & Kajimura, S. Brown and beige fat in humans: thermogenic adipocytes that control energy and glucose homeostasis. The Journal of Clinical Investigation 125, 478-486, doi:10.1172/JCI78362 (2015).

13 Petersen, M. et al. Cardiometabolic characteristics of people with metabolically healthy and unhealthy obesity. Cell Metabolism (2024, [In Press]).

14 Zhang, R. et al. Expression of the Mxra8 Receptor Promotes Alphavirus Infection and Pathogenesis in Mice and Drosophila. Cell Rep 28, 2647-2658 e2645, doi:10.1016/j.celrep.2019.07.105 (2019).

15 Murano, I. et al. Dead adipocytes, detected as crown-like structures, are prevalent in visceral fat depots of genetically obese mice. J Lipid Res 49, 1562-1568, doi:10.1194/jlr.M800019-JLR200 (2008).

16 Cinti, S. et al. Adipocyte death defines macrophage localization and function in adipose tissue of obese mice and humans. J Lipid Res 46, 2347-2355, doi:10.1194/jlr.M500294-JLR200 (2005).

17 Marcelin, G. et al. A PDGFRalpha-Mediated Switch toward CD9(high) Adipocyte Progenitors Controls Obesity-Induced Adipose Tissue Fibrosis. Cell Metab 25, 673-685, doi:10.1016/j.cmet.2017.01.010 (2017).

18 Hepler, C. et al. Identification of functionally distinct fibro-inflammatory and adipogenic stromal subpopulations in visceral adipose tissue of adult mice. Elife 7, doi:10.7554/eLife.39636 (2018).

19 Sun, K., Tordjman, J., Clement, K. & Scherer, P. E. Fibrosis and adipose tissue dysfunction. Cell Metab 18, 470-477, doi:10.1016/j.cmet.2013.06.016 (2013).

20 Sun, K. et al. Endotrophin triggers adipose tissue fibrosis and metabolic dysfunction. Nat Commun 5, 3485, doi:10.1038/ncomms4485 (2014).

21 Chouchani, E. T., Kazak, L. & Spiegelman, B. M. New Advances in Adaptive Thermogenesis: UCP1 and Beyond. Cell Metabolism 29, 27-37, doi:10.1016/j.cmet.2018.11.002 (2019).

22 Brestoff, J. R. et al. Group 2 innate lymphoid cells promote beiging of white adipose tissue and limit obesity. Nature 519, 242-246, doi:10.1038/nature14115 (2015).

23 Cohen, P. et al. Ablation of PRDM16 and beige adipose causes metabolic dysfunction and a subcutaneous to visceral fat switch. Cell 156, 304-316, doi:10.1016/j.cell.2013.12.021 (2014).

24 van Marken Lichtenbelt, W. D. et al. Cold-Activated Brown Adipose Tissue in Healthy Men. New England Journal of Medicine 360, 1500-1508, doi:10.1056/NEJMoa0808718 (2009).

25 Becher, T. et al. Brown adipose tissue is associated with cardiometabolic health. Nature Medicine 27, 58-65, doi:10.1038/s41591-020-1126-7 (2021).

26 Yubero, P. et al. CCAAT/enhancer binding proteins alpha and beta are transcriptional activators of the brown fat uncoupling protein gene promoter. Biochem Biophys Res Commun 198, 653-659, doi:10.1006/bbrc.1994.1095 (1994).

27 Carmona, M. C. et al. Mitochondrial biogenesis and thyroid status maturation in brown fat require CCAAT/enhancer-binding protein alpha. J Biol Chem 277, 21489-21498, doi:10.1074/jbc.M201710200 (2002).

28 Lin, J., Handschin, C. & Spiegelman, B. M. Metabolic control through the PGC-1 family of transcription coactivators. Cell Metabolism 1, 361-370, doi:10.1016/j.cmet.2005.05.004 (2005).

29 Pettersson-Klein, A. T. et al. Small molecule PGC-1α1 protein stabilizers induce adipocyte Ucp1 expression and uncoupled mitochondrial respiration. Molecular Metabolism 9, 28-42, doi:10.1016/j.molmet.2018.01.017 (2018).

30 Feldmann, H. M., Golozoubova, V., Cannon, B. & Nedergaard, J. UCP1 ablation induces obesity and abolishes diet-induced thermogenesis in mice exempt from thermal stress by living at thermoneutrality. Cell Metab 9, 203-209, doi:10.1016/j.cmet.2008.12.014 (2009).

31 Rowland, L. A., Maurya, S. K., Bal, N. C., Kozak, L. & Periasamy, M. Sarcolipin and uncoupling protein 1 play distinct roles in diet-induced thermogenesis and do not compensate for one another. Obesity (Silver Spring) 24, 1430-1433, doi:10.1002/oby.21542 (2016).

32 Zou, W. et al. Myeloid-specific Asxl2 deletion limits diet-induced obesity by regulating energy expenditure. J Clin Invest 130, 2644-2656, doi:10.1172/jci128687 (2020).

33 Sung, H. K. et al. Adipose vascular endothelial growth factor regulates metabolic homeostasis through angiogenesis. Cell Metab 17, 61-72, doi:10.1016/j.cmet.2012.12.010 (2013).

34 Shimizu, I. et al. Vascular rarefaction mediates whitening of brown fat in obesity. J Clin Invest 124, 2099-2112, doi:10.1172/JCI71643 (2014).

35 Xu, Z., Chen, X., Song, L., Yuan, F. & Yan, Y. Matrix Remodeling-Associated Protein 8 as a Novel Indicator Contributing to Glioma Immune Response by Regulating Ferroptosis. Frontiers in Immunology 13, doi:10.3389/fimmu.2022.834595 (2022).

36 Simpson, K. E., Staikos, C. A., Watson, K. L. & Moorehead, R. A. Loss of MXRA8 Delays Mammary Tumor Development and Impairs Metastasis. International Journal of Molecular Sciences 24, 13730 (2023).

37 Han, S. W. et al. DICAM Attenuates Experimental Colitis via Stabilizing Junctional Complex in Mucosal Barrier. Inflamm Bowel Dis 25, 853-861, doi:10.1093/ibd/izy373 (2019).

38 Dieckmann, S. et al. Susceptibility to diet-induced obesity at thermoneutral conditions is independent of UCP1. Am J Physiol Endocrinol Metab 322, E85-E100, doi:10.1152/ajpendo.00278.2021 (2022).

39 Brestoff, J. R. et al. Intercellular Mitochondria Transfer to Macrophages Regulates White Adipose Tissue Homeostasis and Is Impaired in Obesity. Cell Metab 33, 270-282 e278, doi:10.1016/j.cmet.2020.11.008 (2021).

40 Borcherding, N. et al. Dietary lipids inhibit mitochondria transfer to macrophages to divert adipocyte-derived mitochondria into the blood. Cell Metab 34, 1499-1513 e1498, doi:10.1016/j.cmet.2022.08.010 (2022).

