## Extended Data Figures 1-5 for "MXRA8 promotes adipose tissue whitening to drive obesity"

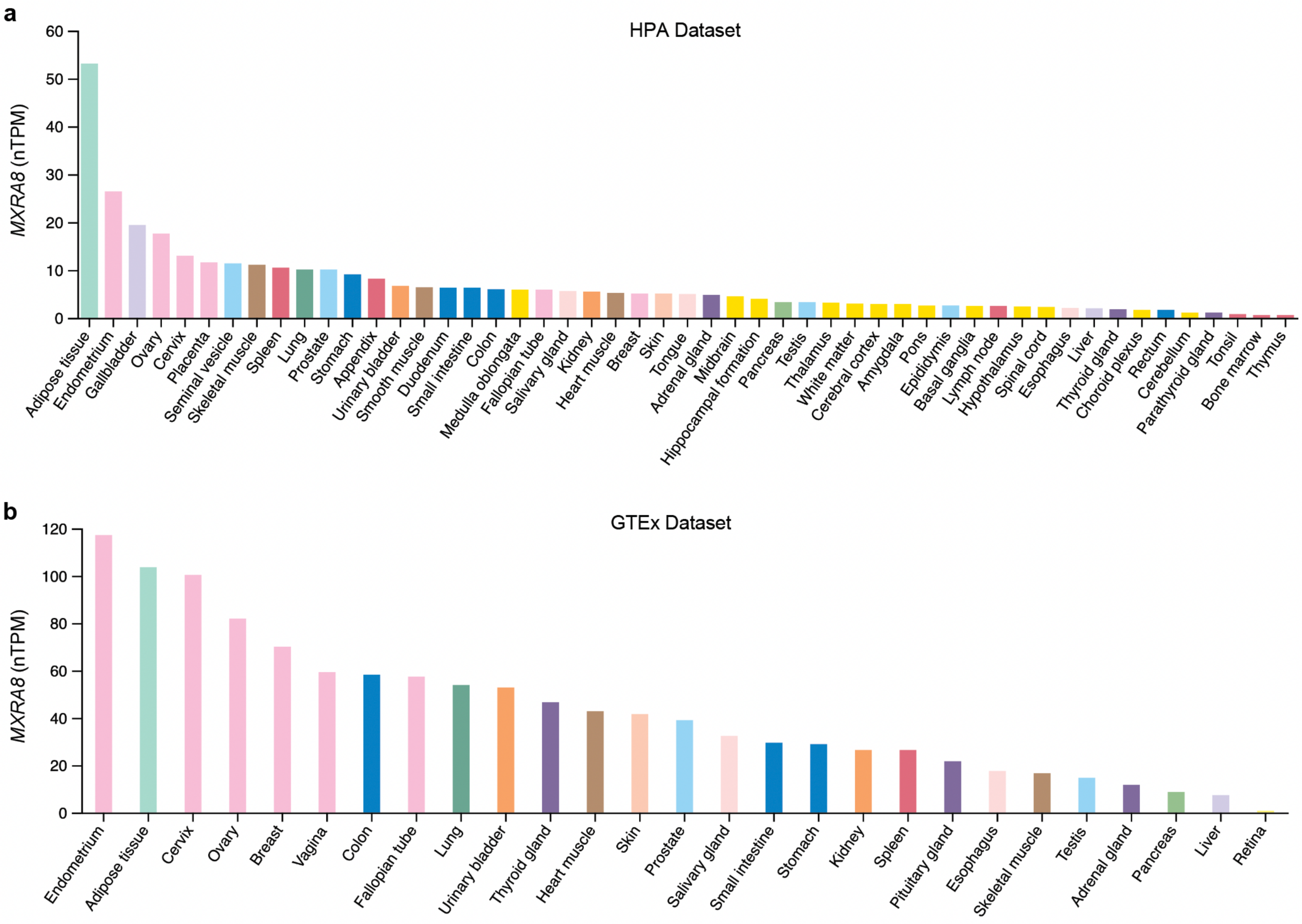


**Extended Data Figure 1. MXRA8 is highly expressed in human adipose tissue. (a)** Human Protein Atlas (HPA) and **(b)** Genotype-Tissue Expression (GTEx) datasets showing *MXRA8* expression in various human tissue organs, rank-ordered based on expression level. These data are related to main Figure 1.

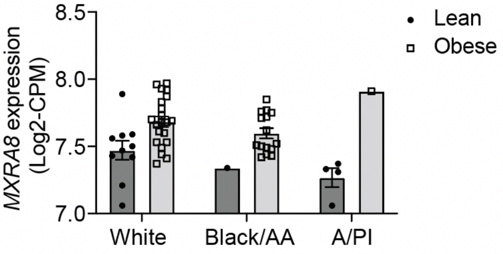


**Extended Data Figure 2. *MXRA8* expression in white adipose tissue is upregulated in human obesity in different ethnicities.** Subcutaneous abdominal adipose tissues were obtained from people who were metabolically healthy lean (MHL, n=15), metabolically healthy obese (MHO, n=18), and metabolically unhealthy obese (MUO, n=19) for bulk RNA-sequencing. The two groups with obesity were combined for this analysis. *MXRA8* expression was compared in the groups who were lean and obese stratified by race: White, Black/African American (AA), and Asian/Pacific Islander (A/PI). These data are related to main Figure 1.

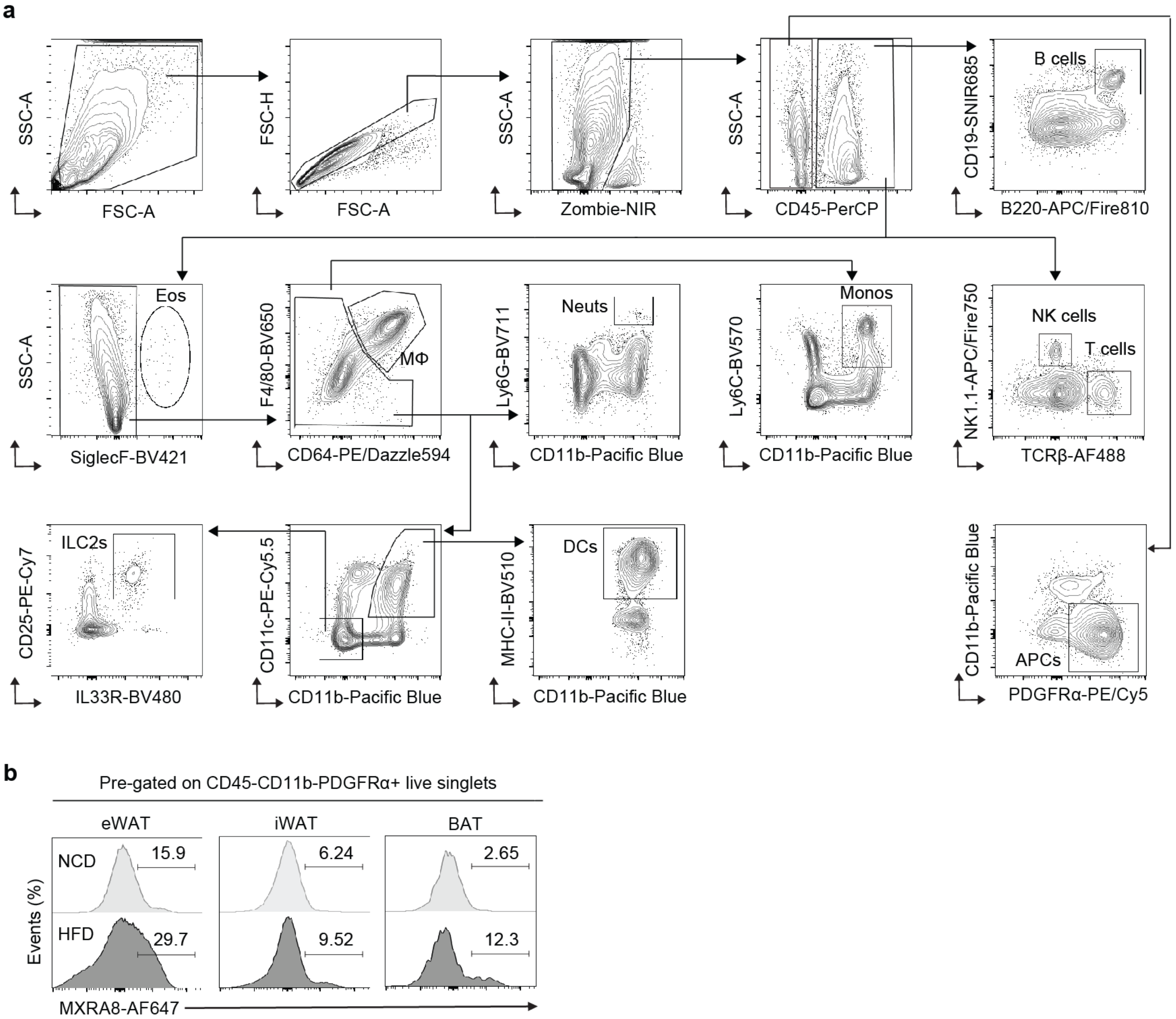


**Extended Data Figure 3. Gating strategy for immune cells and adipocyte progenitor cells and representative flow cytometric plots. (a)** Gating strategy to identify total immune cells, eosinophils (Eos), macrophages (Mφ), neutrophils (Neuts), monocytes (Monos), dendritic cells (DCs), B cells, conventional T cells, natural killer (NK) cells, group 2 innate lymphoid cells (ILC2s), and adipocyte progenitor cells (APs). **(b)** Representative flow cytometry histograms showing surface expression of MXRA8 in APs of eWAT, iWAT, and BAT in wildtype mice fed a normal chow diet (NCD) or high fat diet (HFD, 60% kcal fat) for 10 weeks (age at the start of HFD feeding, 8-12 weeks). These data are related to main Figures 2 and 3.

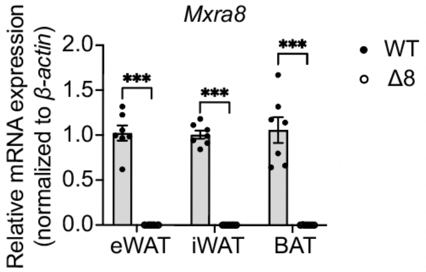


**Extended Data Figure 4. Genotype verification by quantitative RT-PCR.** Relative *Mxra8* mRNA expression in eWAT, iWAT, and BAT of 8-10-week-old male wildtype (WT, n=7) or $\mathit{Mxra8}^{\Delta 8/\Delta 8}$ (Δ8, n=7) mice that were fed a HFD (60% kcal fat) for 12 weeks. Primers were designed to amplify WT *Mxra8* but not the mutant form of the gene. Data are expressed as mean ± standard error of the mean. Two-way ANOVA with Fisher's LSD post hoc test. ***P<0.001. Related to Main figure 3.

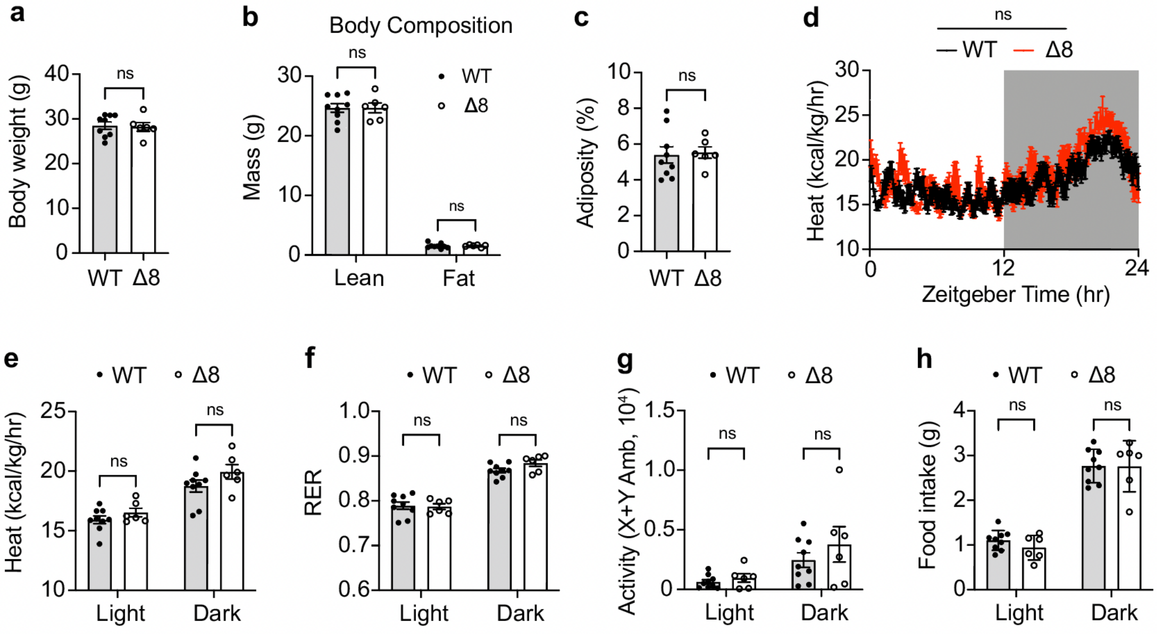


**Extended Data Figure 5. *Mxra8$^{\Delta8/\Delta8}$* and wildtype mice have similar body weights and energy homeostasis on a normal chow diet.** 18-20-week-old male wildtype (WT) (n=9) and *Mxra8$^{\Delta8/\Delta8}$* (Δ8) (n=6) mice were fed a normal chow diet. **(a)** Body weight, **(b)** body composition, and **(c)** adiposity. **(d)** Energy expenditure over a 24h period with the light phase unshaded and the dark phase shaded. **(e)** Average energy expenditure, **(f)** respiratory exchange ratio (RER), **(g)** activity, and **(h)** food intake of WT and Δ8 mice during the light and dark cycles. Data are expressed as mean ± standard error of the mean. For panels a and c, Student's t test. For panels b, two-way ANOVA with LSD post hoc test. For panels d-h, two-way ANOVA with repeated measures and Sidak's post hoc test. These data are related to main Figure 3. Not significant, ns.
